# Protocol for simultaneous evaluation of neuronal activity and neurotransmitter release following chronic amyloid-beta oligomer injections into the hippocampus

**DOI:** 10.1101/2024.09.26.614333

**Authors:** Vincent Hervé, Laurie Bonenfant, Mathilde Amyot, Rime Balafrej, Obai Bin Ka’B Ali, Habib Benali, Jonathan Brouillette

**Author notes:** Technical contact (Vincent Hervé).

## Abstract

In Alzheimer’s disease, there is an imbalance in neurotransmitter release and altered neuronal activation. We present a novel approach to analyze neuronal activity by combining local field potential (LFP) recording with microdialysis within the same animal. This method measures glutamate and GABA levels following chronic hippocampal amyloid-beta oligomer (Aβo) injections in rats. We outline the design of our electrode and canula, the surgical procedure, and the steps for LFP recording, interstitial fluid collection, and Aβo injections simultaneously in living animal.

**Graphical abstract:** 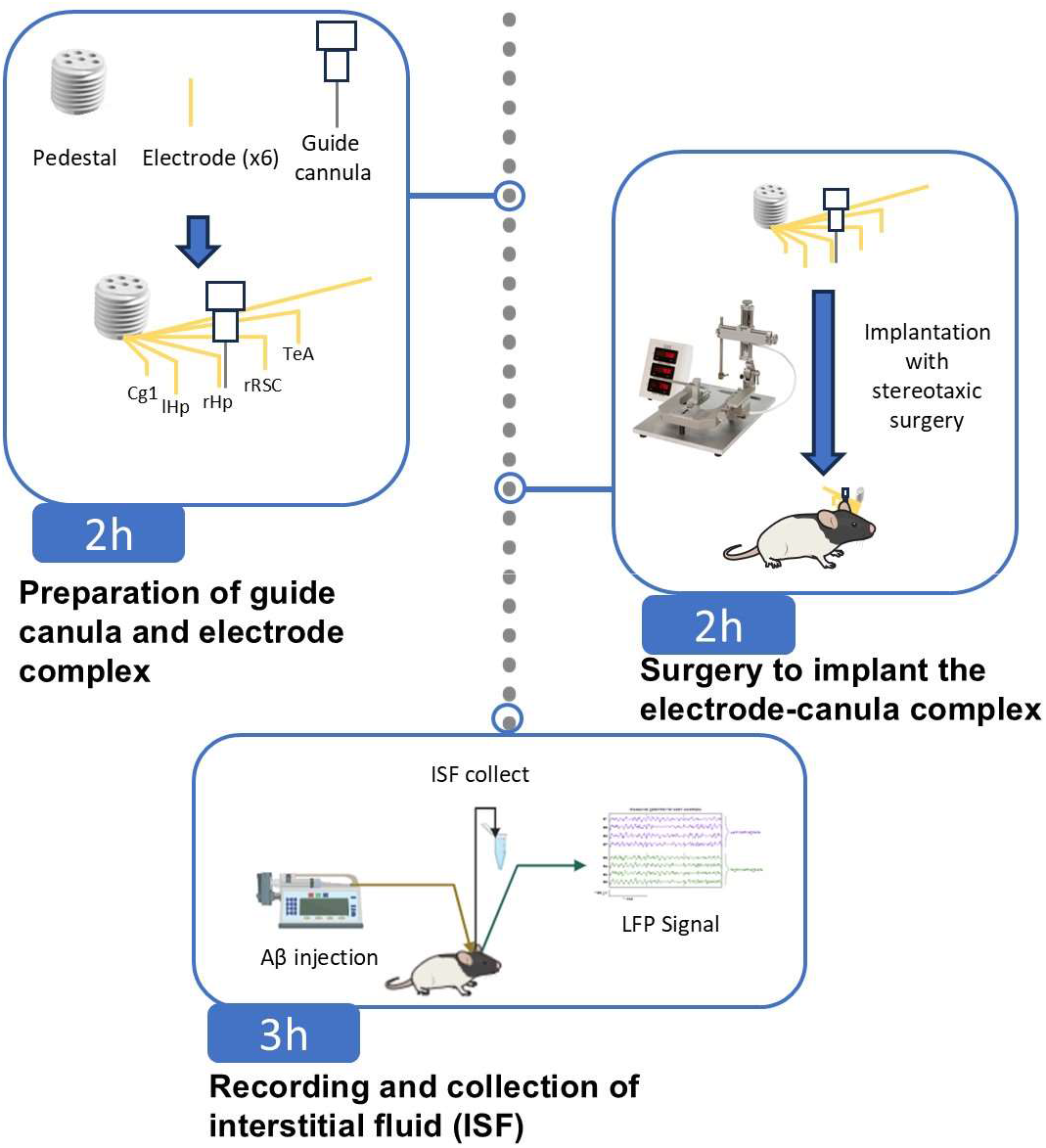

## Before you begin

One of the main neuropathological hallmarks of Alzheimer’s disease (AD) is the accumulation of neurotoxic amyloid-beta oligomers (Aβo), which begins in the brain approximately 15 years prior to the onset of tau pathology, brain atrophy, memory decline, and the clinical diagnosis of AD^1,2^. Thus, by acting on Aβo-induced neurodegeneration we could develop therapies that prevent, or at least slow down, the disease as early as possible before neurodegeneration produces irreversible brain damage and severe cognitive deficits.

Based on animal models, *in vitro* experiments, and human studies, neuronal hyperactivity induced by Aβo has emerged as an early functional characteristic of AD, leading to synaptic deficits, memory dysfunction, and neurodegeneration^3^. Neuronal hyperactivity is a key marker of AD that has been observed in various animal models of the disease as well as in humans^3^. The extracellular increase in endogenous Aβo, induced in part by the inhibition of its degradation, also enhances the release of glutamatergic vesicles and causes neuronal hyperexcitability in primary cultures and rat hippocampal slices^4^. In cultures of neurons derived from induced pluripotent stem cells with certain familial AD mutations that allow for Aβ overexpression, a transient increase in Ca^2+^ and excessive neuronal excitability have been observed^5,6^.

Various studies have elucidated some of the cellular and molecular mechanisms explaining how Aβo can induce neuronal hyperactivity. Indeed, it was suggested that Aβo disrupt the excitation/inhibition balance by reducing inhibitory activity within the GABAergic system, leading to excessive activation of the glutamatergic system in AD mouse models^7-9^. Other studies also suggest that the hyperactivity caused by Aβo may be attributed to excessive accumulation of glutamate in the synaptic cleft due to decreased glutamate reuptake by glial and neuronal cells^10,11^. However, to date, no study has demonstrated *in vivo* if neuronal activity directly associated with Aβo-induced release of glutamate and GABA during the progression of amyloid pathology. Additionally, it remains unclear whether these phenomena are sufficient to lead to excitotoxicity and neuronal death as observed in the hippocampus during the early stages of AD.

The protocol below outlines the various steps required to study neuronal activity and neurotransmitter quantification in the same animal following a chronic injection of Aβo. It describes the procedures for setting up the microdialysis system, performing injections, and recording Local Field Potential (LFP) signals, as well as the surgical steps for electrode and canula implantation. Finally, it details the recording of neuronal signals and the collection of neurotransmitters during Aβo injections. This novel approach is of interest not only to decipher the molecular and cellular mechanism underpinning soluble Aβo, but any other molecules that might have an impact on neurotransmitter release and neuronal activity in different brain regions of animal models.

Before starting this protocol, it is necessary to order the animals, allow them a period of habituation to their housing environment, and accustom them to handling. Additionally, it is necessary to prepare the Aβo solution in advance.

All procedures with animals were conducted according to the guidelines of the Canadian Council on Animal Care, and the protocol was approved by the *Comité de déontologie de l’expérimentation sur les animaux* of the University of Montreal. Laboratories intending to use this protocol must first obtain approval from their respective institutions.

### Institutional permissions

Experiments on live vertebrates must be conducted in accordance with national guidelines and regulations. Additionally, depending on the legislation of certain countries, verify whether the use of anesthetics requires government authorization.

### Animal Habituation

#### 5 days

Rats arrive at the animal facility at 8 weeks of age. The week following their arrival, the animals gradually become accustomed to the experimenters and to being held in restraint. On the first two days, gloved hands are introduced into the cage. On the third day, the animal is placed on a towel held by the experimenter to familiarize it with the scent. From the fourth day until the fifth day, the animal is restrained to simulate an intraperitoneal injection and to mimic the placement of the microdialysis probe.

### Amyloid beta oligomer preparation

#### 3 h

1. Thaw the bottle of Aβ1-42 for 10 min at room temperature then centrifugal at 2500 rpm at 22°C. **Note:** Use the same protocol for the Aβ scramble (control).
2. Add 500 μL of 1,1,1,3,3,3,-Hexafluoro-2-propanol (HFP) under the chemical hood.
  a. Vortex during 1 min 30 sec by inverting the bottle every 30 seconds.
  b. Evaporates the HFP while maintaining a light flow of nitrogen in the bottle.
3. Prepare Tris-EDTA buffer (50 mM Tris, 1 mM EDTA) under sterile conditions:
  a. Add to a 50 mL tube: 2.5ml of 1 M Tris (pH 7.4), 500 µL of 100 mM EDTA, 47 mL sterile water (Baxter JF7624).
  b. Filter this buffer dropwise through a 0.22 µm filter (Sigma Aldricht Z260347-1PAK) using a 20 mL syringe.
4. Add 500 μL of DMSO in the bottle and vortex.
5. Equilibrate the Hitrap desalting column with 25 mL of Tris-EDTA buffer.
  a. Apply 500 μL of Aβ-DMSO in Hitrap desalting column.
  b. Add 1 mL of Tris-EDTA buffer and collect the last 200 μL in tube named Aβ_0_.
  c. Add another 1 mL of Tris-EDTA buffer.
    i. Collect the first 800 μL in low-adhesion tube named Aβ_1_. **Note:** This tube contains the highest concentration of Aβ.
    ii. Collect the last 200 μL in a new tube named Aβ_2_.
6. Determine the concentration of each tube (Aβ_0_, Aβ_1_, Aβ_2_) with Pierce™ BCA Protein Assay Kit. **Note:** Also remember to measure the absorbance of the Tris-EDTA buffer.
  a. Shake the low-adhesion 96-well plate during 10 sec at 120 rpm every 15 min during the 45 min incubation.
  b. Prepare 200 aliquots of 4 μL each from the 800 μL Aβ_1_ solution using low-adhesion tubes and tips. Immediately place the aliquots at -80°C for storage until use.
  c. Measure absorbance of the plate at 562 nm and the concentration in tube Aβ_1_ using the standard line obtained with the absorbances of the different concentrations of BSA measured on the spectrophotometer. **Note:** If the elution of Aβ across the HiTrap desalting column is performed correctly, Aβ should be concentrated entirely in the Aβ_1_ tube, with no presence in the tubes before (Aβ_0_) or after (Aβ_2_) its elution. The concentration in the Aβ_1_ tube should be approximately 0.3 μg/μL.
  d. Determine the volume of Tris-EDTA buffer to add to obtain a final concentration of 0.2μg/μl.

## Key resources table

| REAGENT or RESOURCE | SOURCE | IDENTIFIER |
| --- | --- | --- |
| Chemicals, peptides, and recombinant proteins |  |  |
| Aβ1-42 | rPeptide | #A-1163-2 |
| Alfaxalone (10mg/ml) | Jurox | 424-520425 |
| Dexmedetomidine (0,5mg/ml) | Orion Pharma | 02333929 |
| Meloxicam (5mg/ml) | Vetoquinol | 02487225 |
| Buprénorphine slow release 3mg/ml | Chiron | RxN800431 |
| Atipamazole | Orion Pharma | 02237744 |
| Lidocain | Odan Lab | 02083795 |
| Bioderm | Odan Lab | 00621366 |
| Artificial Cerebrospinal Fluid | Harvard Apparatus | 59-7316 |
| Experimental models: Organisms/strains |  |  |
| Long Evans rats | Charles River | 006L/E |
| Software and algorithms |  |  |
| Harmonie | Natus, Middleton, |  |
| EDF browser | https://www.teuniz.net/edfbrowser/ |  |
| Other |  |  |
| Microdialysis and infusion probe | Basics | MD-2264 |
| Guide Cannula | Basics | MD-2250 |
| 6 Channel Electrode- Platinum Teflon (uncut) | P1 Technologies | E363/6/SPC |
| 6 Channel Electrode Pedestal | P1 Technologies | MS363 |

## Step-by-step method details

### Preparation of guide canula and electrode complex

#### 2 h

**This step involves assembling the electrode complex and guide cannula to prepare them for implantation in the animal**.

1. Ensure that the stylet of the cap is the same length as the microdialysis probe. Place the cap in the guide canula and then into the canula holder (Fig. 1A, item 3; Kopf instruments - Model 1766-AP). **Note**: If the cap is too large to fit into the canula holder, it can be sanded down beforehand.
2. Place the canula holder with the guide canula perpendicular to the stereotaxic table, with the canula facing upwards.
3. Place a cut needle at the end of the standard electrode holder (Fig. 1A, item 2; Kopf instruments - Model 1770) which is attached directly to the stereotaxic apparatus (Fig. 1A, item 1) to adjust the precise coordinates.
4. Place each electrode in the 6-channel electrode pedestal. **Critical:** Electrodes are very fragile when connected to the pedestal.
5. Place the cap on the pedestal and wrap it with sturdy scotch tape (in red **Figure 1**) to create an extra thickness of approximately 2 mm. **Note:** This step is important so that the animal can be connected without obstruction during EEG recordings.
6. Wrap the pedestal with scotch tape to fix it with the guide canula.
7. Place and adjust all the electrodes using the stereotaxic apparatus according to the tip of the stylet inserted into the guide canula. This serves as the “0” reference point for the coordinates (mediolateral (ML), anteroposterior (AP), dorsoventral (DV)) for positioning each electrode.
8. Using the needle of the standard electrode holder attached to the stereotaxic apparatus, adjust each electrode according to the following coordinates:
9. Trim the electrodes to match the specified coordinates, and carefully scrape off less than 1 mm from the tip of each electrode to remove the insulating layer. Apply a few drops of dental cement to the electrode side of the pedestal to secure the joint between the electrodes and the pedestal.

**Figure 1.**
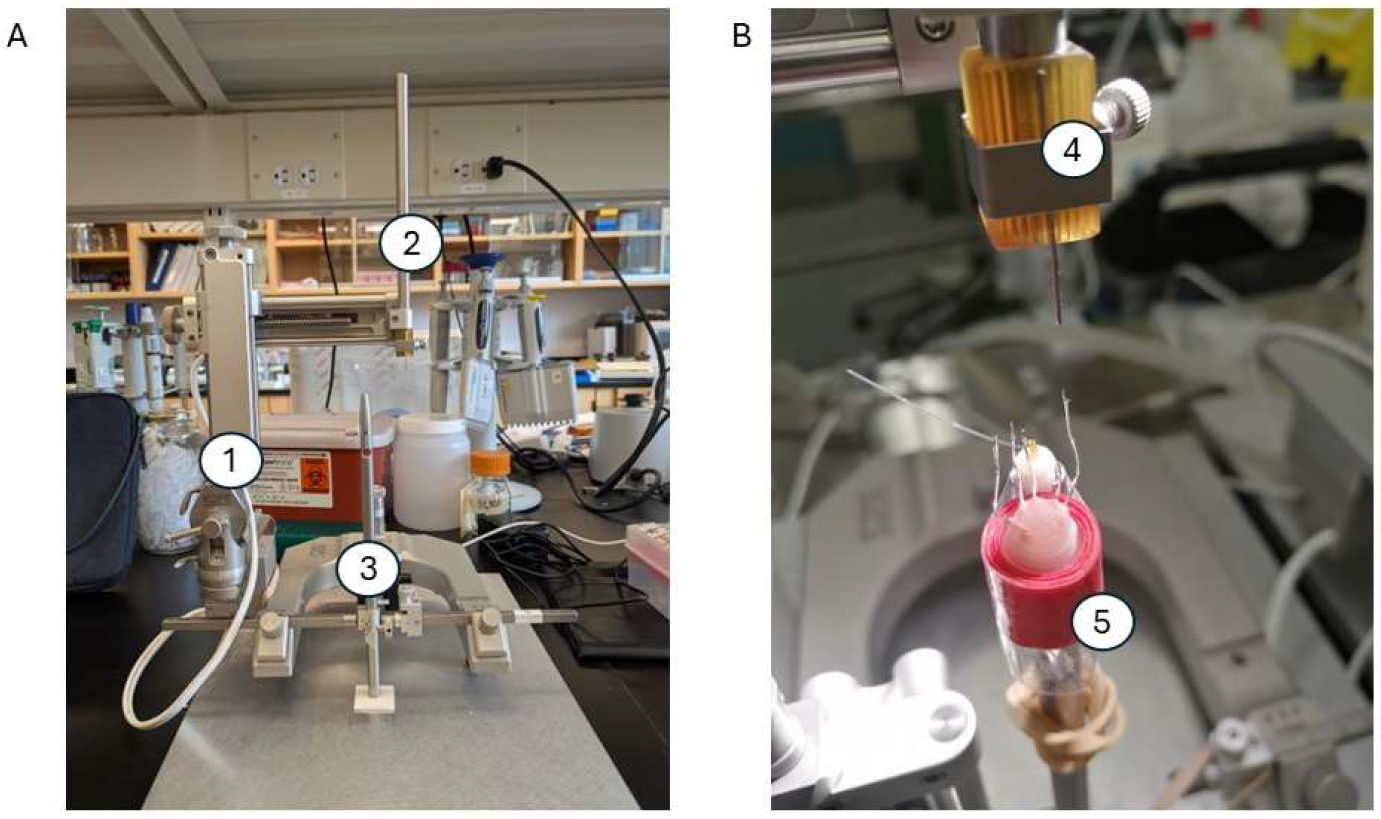
Preparation of guide canula and electrode complex. (A) Setup for assembling the electrodes and guide canula using a stereotaxic apparatus (1). The standard electrode holder with cut needle (2) and canula holder (3) are shown. (B) The pedestal containing the five electrodes and the guide canula (5) is shown, along with the cut needle on the standard electrode holder.

**Table 1:** Coordinates of the different regions where the electrodes are implanted, with the guide canula and its electrode (right hippocampus) as the reference point.

| (mm) | Right Hippocampus (rHp) | Left Hippocampus (lHp) | Right RSC | Right TeA | Right Prefrontal cortex (Cg1, Cg2) |
| --- | --- | --- | --- | --- | --- |
| ML | 0,00 | -3,20 | -0,80 | 5,44 | -1,00 |
| AP | 0,00 | 0,00 | 2,00 | 2,20 | -5,50 |
| DV | 0,00 | 0,00 | 1,00 | -2,40 | 0,60 |

### Surgery to implant the electrode-canula complex

#### 2h

10. Surgeries are performed in a sterile environment with the use of gloves, a mask, and a hair bonnet during the procedure. All surgical tools are pre-cleaned and sterilized by autoclaving. At 10 weeks of age, the rats undergo surgery to implant the electrodes and the guide canula for the microdialysis probe.
11. Administer a subcutaneous injection of slow-release buprenorphine (72-hour formulation) at least 5 h prior to surgery.
12. Anesthetize the rat with an intraperitoneal injection of a cocktail containing alfaxalone (20 mg/kg) and dexmedetomidine (0.1 mg/kg). **Note:** Ensure the absence of reflexes before proceeding.
  a. Inject Meloxicam 1mg/kg subcutaneously.
  b. Shave the animal at the incision site with clippers before placing it on the stereotaxic frame.
  c. Place the animal on the stereotaxic apparatus using ear bars to immobilize the head. The animal is placed on a heating mat maintained at 37°C.
  d. Position the mask over the rat’s nose to maintain anesthesia with isoflurane at a flow rate of 0.5 to 1.0%. **Critical:** After fitting the mask and ear bars, ensure the head remains stable when pressure is applied.
  e. Apply lubricating eye ointment during anesthesia to prevent dryness.
  f. Sterilize the incision area with an alcohol swab.
  g. Make an incision in the skin above the skull.
  h. Locate the bregma and lambda, and mark them with a felt-tip pen. Here, the bregma serves as the “0” reference point for the coordinates on the stereotaxic device. To ensure the skull is flat, a difference of 0.05 cm or less in the dorsoventral axis between the bregma and the lambda is considered acceptable.
13. Mark each point corresponding to the electrode site as follow:
  a. Drill a hole for each electrode using a 1/32’’ drill bit, and use a 3/64’’ drill bit for the guide canula and the adjacent electrode. Additionally, make two more holes with a 1/32’’ drill bit to secure the mounting with two screws placed on either side of the bregma.
  b. Add a hole for the gold screw that serves as the reference electrode. The screw is placed above the cerebellum, which is known to be a control zone. **Note**: The cerebellum is highly vascularized, so if hemorrhaging occurs, wait for it to stop before placing the screw.
  c. On the stereotaxic frame, replace the standard electrode holder containing the needle with the canula holder containing the electrode and canula assembly.
  d. Adjust the arm with the canula holder so that the electrodes align directly with the drill holes.
  e. A solder wire is wrapped around and soldered to the gold-plated screw (the reference electrode) located above the cerebellum. **Note**: The cerebellum is an ideal control structure for recording reference signals in LFP measurements.
14. Acrylic dental cement is then applied around the guide canula, the electrode pedestal, and the screws to secure the setup. **Note:** Remove the tape binding the pedestal and the guide cannula to improve accessibility and facilitate better fixation of the assembly.
15. **Critical:** Apply multiple layers of dental cement to ensure the assembly is firmly secured and to prevent any movement.
16. Sutures are applied to close the skin using 4-0 surgical thread. Trim the rat’s nails once the sutures are complete.
  a. Administer 3 ml of saline subcutaneously at 37°C. Apply a cream mixture of lidocaine and polysporin to the incision site daily for 2 days following the surgery.
  b. Inject Atipamazole at an equivalent dose (v/v) of Dexmedetomidine to induce rapid awakening of the animal. Place the rat in an oxygen chamber heated to 37°C.
  c. Once the rat is awake and able to walk, return it to its cage and housing room.

**Table 2:** Stereotaxic coordinates of the different brain regions where the skull holes are made for the implantation of our electrode and canula system. RSC = retrosplenial cortex, TeA = temporal association cortex, and PFC = prefrontal cortex.

| (mm) | Right Hippocampus | Left Hippocampus | Right RSC | Right TeA | Right Prefrontal cortex | Reference Electrode |
| --- | --- | --- | --- | --- | --- | --- |
| ML | 1,60 | -1,60 | 0,80 | 7,04 | 0,60 | -0,94 |
| AP | 3,80 | 3,80 | 5,80 | 6,00 | -1,70 | 10,50 |
| DV | -3,60 | -3,60 | -2,60 | -6,00 | -3,00 | N/A |

**Figure 2.**
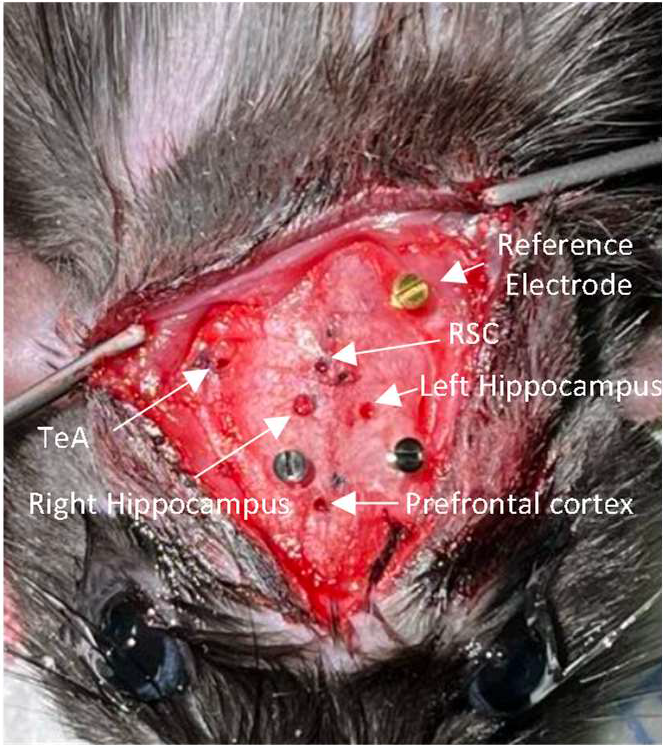
Image depicting the holes made in the animal’s skull and the positions of the three screws.

### Recording and collection of interstitial fluid (ISF)

#### 3h

17. At 12 weeks of age, bring the rats into the LFP recording room and allow them a 30-min habituation period.
18. During this 30-min period, prepare the setup for microdialysis:
  a. Take 1 mL of artificial cerobrospinal fluid (CSF) (Harvard Apparatus) with a 3 mL syringe.
  b. Place the syringe on a pump (CMA pump).
  c. Cut the needle 23G to remove the pointed tip, and connect the syringe to a swivel (Instech 375/D/25) with PE50 tubing.
  d. Connect the swivel to the microdialysis probe (yellow channel).
  e. Connect the microdialysis probe (green channel) to the swivel with microtubing (Basics MD-1511).
  f. Calculate the volume between the microdialysis outlet and the fraction collection to adjust the timing of collection with LFP recording.
19. Set the pump flow rate to 1 µL/min and ensure that the CSF is circulating properly by observing the appearance of a drop at the end of the circuit.
20. Place the rat in a new cage and position the microdialysis probe on the animal. Allow a 1-hour equilibration period for the microdialysis probe. During this period, check the volume of the fractions multiple times to ensure 10 µl per 10 minutes. **Critical:** Monitor the animal to prevent it from biting the tubes, which could cause leaks and require repeating the experiment.
  a. After calculating the volume between the microdialysis probe outlet and the collection (7 µL), connect the animal for LFP recording 7 min before starting the ISF fraction collection. **Note:** Keep the animal awake throughout the experiment.
  b. After 1 h of equilibration, collect fractions every 10 min, each with a volume of 10 μL. Add 1 µL of 0.25 mol/L perchloric acid to each fraction to prevent analyte degradation.
  c. After 30 min of microdialysis, perform an injection of Aβo (preparation described in “Before You Begin”) at a concentration of 0.2 µg/µl using a microsyringe (10 µL) pump at a flow rate of 1 µL/min for 2 minutes. The injection channel is the red channel in the microdialysis probe. Keep the Aβo solution at room temperature for 1 h before injection to allow the oligomerization of Aβ.
  d. Allow 8 minutes after the injection before removing the injection channel.
21. The LFP signals are amplified using a Lamont amplifier, sampled at 1024 Hz, and filtered with the commercial software Harmonie (Natus, Middleton, WI, USA) as previously described^12^.
22. After the experiment, disconnect the animal from the microdialysis and LFP channels, and return it to its cage.
  a. Clean the probe by flushing the injection and microdialysis channels with water.
  b. Connect the green and yellow channels to PE-50 tubing filled with water, and plug the red channel. Leave probe to soak in water until next use. **Note:** Store the probe in water within its cap at 4°C.
  c. Clean the swivel with a few microliters of water, then dry it with a connected syringe at a flow rate of 10 μl/min.

**Figure 3.**
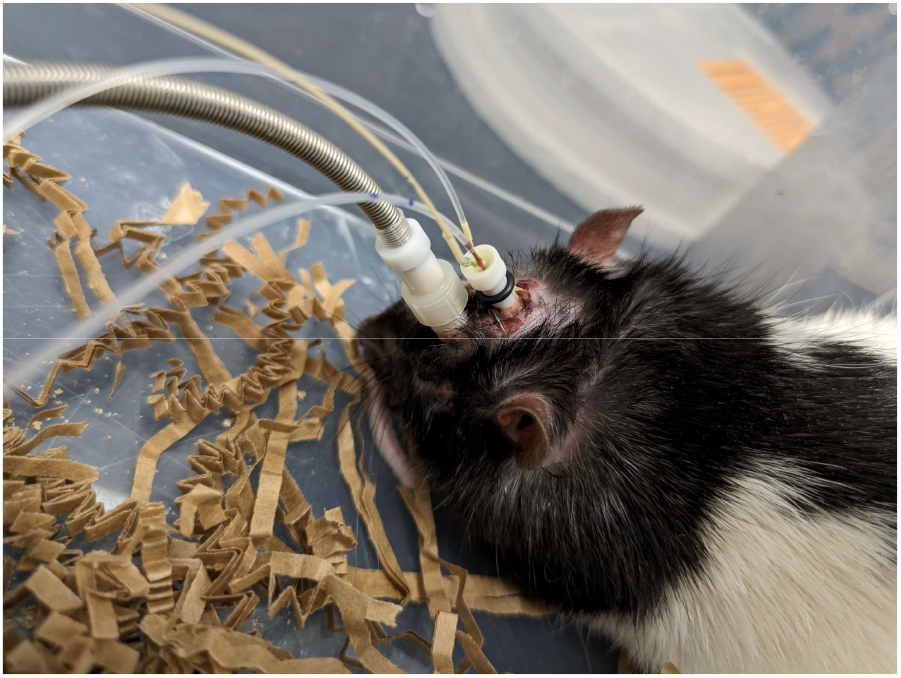
Image of a rat with the microdialysis probe and connected for LFP recording.

### Expected outcomes

Using the novel animal model described here, we observed that soluble Aβo injections over 5 days in the right hippocampus of rats induced a clear time-dependent increase in delta power (slope = 0.73, SE = 0.13, *p* < 0.001) and a progressive decrease in higher frequencies, including theta (slope = -0.52, SE = 0.22, *p* = 0.018), alpha (slope = -0.58, SE = 0.16, *p* < 0.001), and beta (slope = -0.57, SE = 0.21, *p* = 0.006), in the injected hippocampus (Fig. 4a). Gamma power also shows a decrease over time (slope = -0.62, SE = 0.37, *p* = 0.094), though this effect approaches marginal significance. This pattern of increased slow-frequency and decreased higher-frequency power reflects the slowing of neuronal oscillations commonly associated with AD pathology. These results are in accordance with what we observed in a previous study^13^, confirming that Aβo injections consistently lead to these time-dependent changes in power. In the left hippocampus with no injection (Fig. 4c), the Aβo group initially showed stable trends, with a late emergence of increased delta power and decreased higher-frequency power on day 5, resembling those observed in panel (a). This suggests a delayed or propagating effect from the right hippocampus. In contrast, the Aβscramble (Aβscr) control group showed no significant changes in power over time, with no substantial deviations from baseline (Fig. 4b,d).

**Figure 4.**
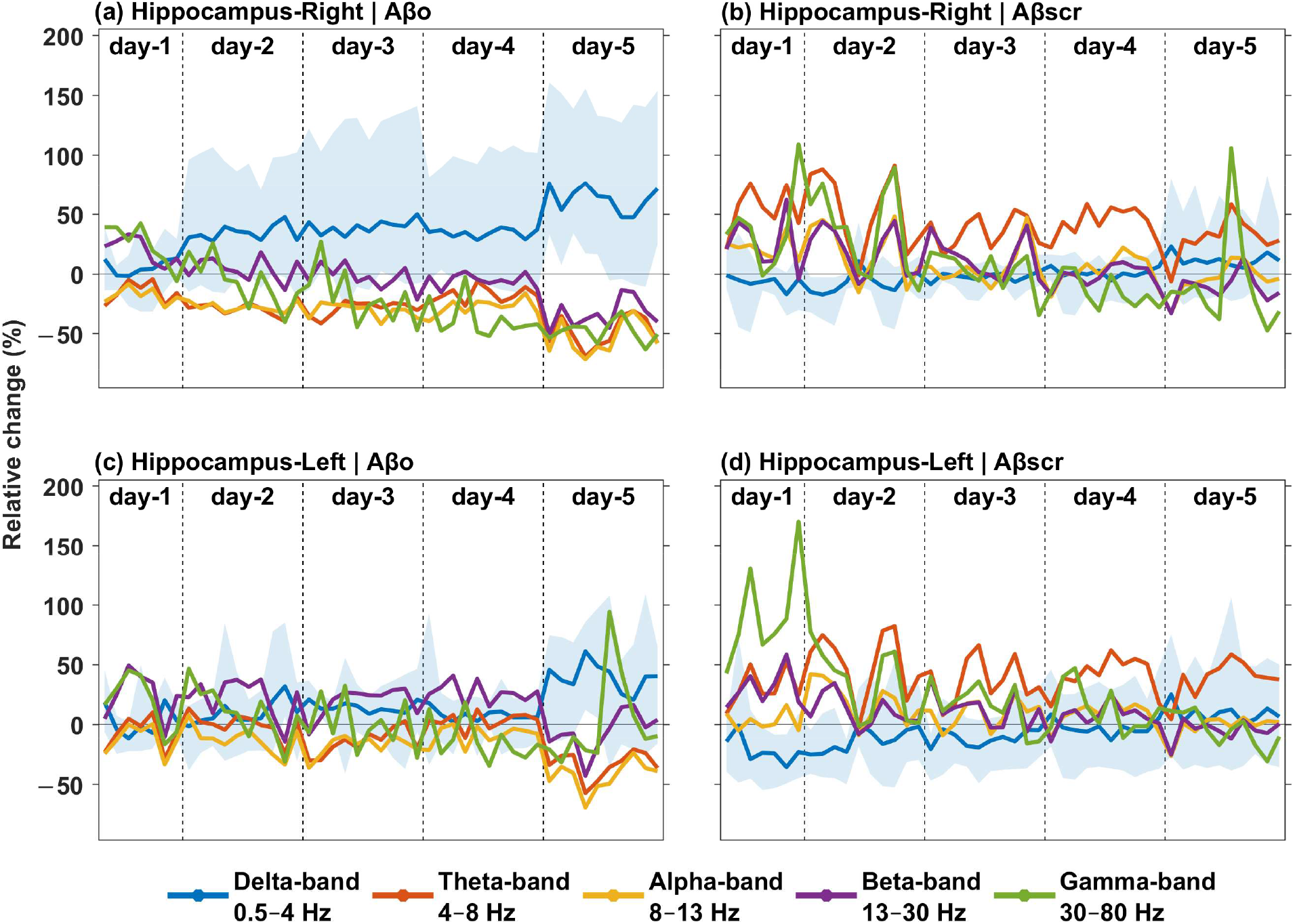
Effects of chronic Aβo injections over 5 days on LFP power across canonical frequency bands, plotted as the relative change in power (%) from baseline over time. Comparisons were made between the right hippocampus (injection site) and the left hippocampus for both the Aβo group (n = 4) and the Aβscr control group (n = 5). A semi-transparent ribbon is overlaid on the delta band to show the minimum and maximum values, calculated across subjects. Significance was assessed at a threshold of *p* < 0.05. Statistical trends were analyzed using linear mixed-effects models for each frequency band, with fixed effects for time, group, and hemisphere, and random intercepts for subject-level variability.

In future studies, this animal model could allow for the investigation of the impact of hippocampal Aβo injections on other brain regions, particularly the default-mode network (DMN), where functional connectivity disruptions are detected in early AD^14^. The DMN is involved in several high-level cognitive processes such as semantic memory, social cognition, and decision-making^15^, and represents a network of brain regions that are typically more active during rest or when attention is not focused on the external environment^16^

By implanting three additional electrodes in different areas of the DMN (PFC, RSC, TeA), this new animal model enables the assessment of whether Aβo accumulation in the hippocampus affects its interactions with this neural network. Specifically, this approach allows for the analysis of frequency band profiles and functional connectivity measures (e.g., coherence, phase-amplitude coupling) between the hippocampus and DMN regions across different time points. This will help determine whether similar network-wide changes occur alongside those observed in the hippocampus during the injection period. This in vivo model can also be used to explore whether changes in neuronal dynamics are directly related to glutamate and GABA release levels following Aβo injection into the hippocampus. Using another group of rats, neurodegeneration, along with astrocytic and microglial activation, can be assessed via immunofluorescence using antibodies targeting NeuN (neuronal marker), GFAP (astrocyte marker), and IBA1 (microglia marker) proteins after different Aβo and Aβscr injection intervals. This will allow for the evaluation of neuronal death, astrogliosis, and microgliosis in brain regions of interest, potentially linking these molecular and cellular changes to the observed alterations in neuronal activity and connectivity.

### Troubleshooting

#### Problem 1

The electrode pedestal is detached from the animal after the first day of LFP recording.

#### Potential solution

We encountered this issue with our first batch of animals and addressed it by increasing the number of screws on the animal’s skull from one to three. Since then, the problem has not recurred. Additionally, during the surgery, ensure that the pedestal and guide canula are thoroughly coated with dental cement to secure the assembly. The screws will aid in this fixation. Make sure to wait until the dental cement has fully solidified before proceeding.

#### Problem 2

The collection volume is not 10 µL or no fluid is emerging from the tube after microdialysis.

#### Potential solution

- Check all the tubing connections to ensure there are no leaks at their junctions, and inspect the wires, as the rat may have bitten one. To prevent this, carefully monitor the animal and use shorter tubing to minimize accessibility and reduce the risk of biting.
- If the microdialysis probe is blocked, restart the experiment with the probe balanced if the injection has not yet occurred. If a tube is chewed, replace the tubing and restart the experiment. Ensure the probe is not blocked in either the microdialysis or injection channels before placing it on the animal.

## Acknowledgments

The authors are thankful to Ian Massé and Louis De Beaumont for help and advice to setup the microdialysis procedure. The authors also want to thank Louiza Mahrouche and Mihaela Friciu from the Platform of Biopharmacy at the University of Montreal for quantifying neurotransmitters by high-performance liquid chromatography coupled with mass spectrometry.

The research has been funded by a grant from the Canadian Institutes of Health Research to J.B., a scholarship from the Fonds de recherche du Québec - Santé to V.H., and a scholarship from the PRogramme d’Excellence en Médecine pour l’Initiation En Recherche to R.B. and L.B.. H.B. and O.B.K.A. are funded by a Canada Research Chair from the Natural Sciences and Engineering Research Council of Canada (grant number NC0981).

## Author contributions

Conceptualization and methodology, V.H, L.B.; M.A., R.B., J.B.; Investigation, V.H, J.B.; Data analysis O.B.K.A, H.B.; Writing, V.H, J.B; Funding acquisition, supervision, and project administration, J.B. All authors approved the final version of the manuscript.

## References

1. Cline, E.N., Bicca, M.A., Viola, K.L., and Klein, W.L. (2018). The Amyloid-beta Oligomer Hypothesis: Beginning of the Third Decade. J Alzheimers Dis 64, S567–S610. 10.3233/JAD-179941.

2. Brouillette, J. (2014). The effects of soluble Abeta oligomers on neurodegeneration in Alzheimer’s disease. Curr Pharm Des 20, 2506–2519. 10.2174/13816128113199990498.

3. Hector, A., and Brouillette, J. (2020). Hyperactivity Induced by Soluble Amyloid-beta Oligomers in the Early Stages of Alzheimer’s Disease. Front Mol Neurosci 13, 600084. 10.3389/fnmol.2020.600084.

4. Abramov, E., Dolev, I., Fogel, H., Ciccotosto, G.D., Ruff, E., and Slutsky, I. (2009). Amyloid-beta as a positive endogenous regulator of release probability at hippocampal synapses. Nat Neurosci 12, 1567–1576. 10.1038/nn.2433.

5. Ghatak, S., Dolatabadi, N., Trudler, D., Zhang, X., Wu, Y., Mohata, M., Ambasudhan, R., Talantova, M., and Lipton, S.A. (2019). Mechanisms of hyperexcitability in Alzheimer’s disease hiPSC-derived neurons and cerebral organoids vs isogenic controls. Elife 8. 10.7554/eLife.50333.

6. Park, J., Wetzel, I., Marriott, I., Dreau, D., D’Avanzo, C., Kim, D.Y., Tanzi, R.E., and Cho, H. (2018). A 3D human triculture system modeling neurodegeneration and neuroinflammation in Alzheimer’s disease. Nat Neurosci 21, 941–951. 10.1038/s41593-018-0175-4.

7. Busche, M.A., Eichhoff, G., Adelsberger, H., Abramowski, D., Wiederhold, K.H., Haass, C., Staufenbiel, M., Konnerth, A., and Garaschuk, O. (2008). Clusters of hyperactive neurons near amyloid plaques in a mouse model of Alzheimer’s disease. Science 321, 1686–1689. 10.1126/science.1162844.

8. Busche, M.A., and Konnerth, A. (2016). Impairments of neural circuit function in Alzheimer’s disease. Philos Trans R Soc Lond B Biol Sci 371. 10.1098/rstb.2015.0429.

9. Palop, J.J., and Mucke, L. (2010). Amyloid-beta-induced neuronal dysfunction in Alzheimer’s disease: from synapses toward neural networks. Nat Neurosci 13, 812–818. 10.1038/nn.2583.

10. Harris, M.E., Wang, Y., Pedigo, N.W., Jr., Hensley, K., Butterfield, D.A., and Carney, J.M. (1996). Amyloid beta peptide (25-35) inhibits Na+-dependent glutamate uptake in rat hippocampal astrocyte cultures. J Neurochem 67, 277–286. 10.1046/j.1471-4159.1996.67010277.x.

11. Parpura-Gill, A., Beitz, D., and Uemura, E. (1997). The inhibitory effects of beta-amyloid on glutamate and glucose uptakes by cultured astrocytes. Brain Res 754, 65–71. 10.1016/s0006-8993(97)00043-7.

12. Seok, B.S., Cao, F., Belanger-Nelson, E., Provost, C., Gibbs, S., Jia, Z., and Mongrain, V. (2018). The effect of Neuroligin-2 absence on sleep architecture and electroencephalographic activity in mice. Mol Brain 11, 52. 10.1186/s13041-018-0394-3.

13. Hector, A., Provost, C., Delignat-Lavaud, B., Bouamira, K., Menaouar, C.A., Mongrain, V., and Brouillette, J. (2023). Hippocampal injections of soluble amyloid-beta oligomers alter electroencephalographic activity during wake and slow-wave sleep in rats. Alzheimers Res Ther 15, 174. 10.1186/s13195-023-01316-4.

14. Warren, J.D., Rohrer, J.D., Schott, J.M., Fox, N.C., Hardy, J., and Rossor, M.N. (2013). Molecular nexopathies: a new paradigm of neurodegenerative disease. Trends Neurosci 36, 561–569. 10.1016/j.tins.2013.06.007.

15. Mars, R.B., Neubert, F.X., Noonan, M.P., Sallet, J., Toni, I., and Rushworth, M.F. (2012). On the relationship between the “default mode network” and the “social brain”. Front Hum Neurosci 6, 189. 10.3389/fnhum.2012.00189.

16. Shulman, G.L., Fiez, J.A., Corbetta, M., Buckner, R.L., Miezin, F.M., Raichle, M.E., and Petersen, S.E. (1997). Common Blood Flow Changes across Visual Tasks: II. Decreases in Cerebral Cortex. J Cogn Neurosci 9, 648–663. 10.1162/jocn.1997.9.5.648.

